# Progerin-Expressing Endothelial Cells are Unable to Adapt to Shear Stress

**DOI:** 10.1101/2021.08.20.456996

**Authors:** Brooke E. Danielsson, Hannah C. Peters, Kranthi Bathula, Lindsay M. Spear, Natalie A. Noll, Kris N. Dahl, Daniel E. Conway

## Abstract

Hutchinson-Gilford Progeria Syndrome (HGPS) is a rare premature aging disease caused by a single-point mutation in the lamin A gene, resulting in a truncated and farnesylated form of lamin A. This mutant lamin A protein, known as progerin, accumulates at the periphery of the nuclear lamina, resulting in both an abnormal nuclear morphology and nuclear stiffening. HGPS patients experience rapid onset of atherosclerosis, with death from heart attack or stroke as teenagers. Progerin expression has been shown to cause dysfunction in both vascular smooth muscle cells and endothelial cells (ECs). In this study we examined how progerin-expressing ECs adapt to fluid shear stress, the principal mechanical force from blood flow. We compared the response to shear stress for progerin-expressing, wild-type lamin A overexpressing, and control ECs to physiological levels of fluid shear stress. Additionally, we also knocked down ZMPSTE24 in ECs, which results in increased farnesylation of lamin A and similar phenotypes to HGPS. Our results showed that ECs either expressing progerin or with ZMPSTE24 knockdown were unable to adapt to shear stress, experiencing significant cell loss at a longer duration of exposure to shear stress (3 days). ECs overexpressing wild-type lamin A also exhibited similar impairments in adaptation to shear stress, including similar levels of cell loss. Quantification of nuclear morphology showed that progerin-expressing ECs had similar nuclear abnormalities in both static and shear conditions. Treatment of progerin-expressing cells and ZMPSTE24 KD cells with lonafarnib and methystat, drugs previously shown to improve HGPS nuclear morphology, resulted in improvements in adaptation to shear stress. Additionally, pre-alignment of cells to shear stress prior to progerin-expression prevented cell loss. Our results demonstrate that changes in nuclear lamins can affect the ability of EC to properly adapt to shear stress.

## Introduction

The nuclear lamina, a fibrous lamin-protein network, located between the inner membrane of the nuclear envelope and chromatin, provides structural support for the nucleus and plays a major role in nuclear shape, gene regulation, as well as the assembly/disassembly of the nucleus during cell division (1–4). Numerous studies have shown the mechanoadapative and mechanoresponsive nature of the nuclear lamina under force (5–9).

Hutchinson-Gilford progeria syndrome (HGPS) is a rare premature aging disease in children caused by an autosomal dominant mutation in the *LMNA* gene, resulting in an aberrant form of lamin A termed progerin(10). The predominant mutation in HGPS involves a de novo point mutation (1824C>T), which activates a cryptic donor splice site resulting in an internal deletion of 50 amino acids. These amino acids include a cleavage site for ZMPSTE24, a protease that removes the farnesyl group from mature lamin A—loss of this cleavage site leads to the permeant farnesylation of progerin. Permeant farnesylation of progerin is thought to be the critical feature of HGPS, which is supported by the observation that ZMPSTE24 knockout mice phenocopy HGPS (11, 12). Similar to lamin A, progerin accumulates at the nuclear periphery. However, progerin expression has been shown to alter the nuclear lamina structure, and leads to several downstream nuclear defects, including: abnormal nuclear morphology, increased nuclear stiffness, redistribution of heterochromatin, modified nuclear pore structure, alterations in gene expression, and nuclear structural instability (13–17).

HGPS patients have accelerated atherosclerosis, leading to premature death as a result of heart attack and stroke (18). Cardiovascular alterations in HGPS patients are similar to atherosclerosis of aging individuals such as exhibiting hypertension, vascular stiffening and calcification, and plaque formation(19–21). Yet these patients do not experience traditional risk factors for atherosclerosis, such as hypercholesterolemia or increased serum levels of creactive protein(22). Thus, an important question is why and how progerin expression affects the vasculature to lead to rapid onset of atherosclerosis? One of the hallmarks of the disease is the loss of vascular smooth muscle cells (vSMCs) in large arterial vessels(23, 24). Several studies, using both iPSCs and HGPS mouse models have shown that expression of progerin in vSMCs impairs cell proliferation (25), impairs cell response to physiological levels of strain (stretch) (26, 27), reduces vasoreactivity (28), and accelerates atherosclerosis by inducing endoplasmic reticulum stress (29, 30).

In addition to vSMCs, progerin also alters endothelial cell function. Using a HGPS mouse model with progerin expression only in endothelial cells it was shown that this resulted in increased inflammation, impaired vascularization, and shortened life span(31). In a similar HGPS mouse endothelial model, it was observed that progerin-expressing endothelial cells caused cardiac pathologies, and that these endothelial cells had impaired mechanoresponsivity(32). We hypothesized that an important aspect of progerin expression in endothelia cells would be impaired mechanoadaptation to shear stress, which was suggested in a previous study(32).To investigate this hypothesis, we developed an HGPS model using human umbilical vein endothelial cells (HVUEC) expressing progerin, as well as HUVECs with ZMPSTE24 shRNA knockdown. Progerin-expressing cells, as well as ZMPSTE24 knockdown cells, failed to adapt to physiological levels of fluid shear stress, exhibiting cell loss at longer timepoints of shear stress exposure. Cell loss was rescued by treatment with the farnesyltranferase inhibitor lonafarnib, DNA demethylase inhibitor methylstat, as well as preadaptation of cells to fluid shear stress prior to progerin expression. Collectively our results show that endothelial cells expressing progerin cannot adapt to the mechanical forces of fluid shear stress, which may be an important aspect of the rapid onset of atherosclerosis in HGPS patients.

## Materials and Methods

### Cell Culture and Transfection

Commercially available primary human umbilical vein endothelial cells (HUVECs) (pooled, passages 3-5, Lonza, Basel, Switzerland) were grown in EGM-2 medium (Lonza, Basel, Switzerland). To express progerin in HUVECs, a previously described HA-tagged progerin adenovirus was developed(33).The lowest level of adenovirus that infected nearly 100% of cells was used. To overexpress wild-type lamin A in HUVECs, lamin A adenovirus (based on RefSeq BC014507) was purchased from Vector Biolabs and used at an identical titer level as progerin, as previously described(34).

### shRNA Knockdown

To knockdown ZMPSTE24 the pLKO-1 vector was used (Sigma Aldrich, clone ID TRCN0000294124, target sequence TGGTAAGGCCAATGTTATTTA). Lentivirus was prepared using HEK 293 cells, with second generation packaging plasmids, pSPAX2 (Addgene 12260) and pMD2.G (Addgene 12259). HUVECs were transduced with lentivirus shRNA and selected with puromycin (1 μg/ml). A non-targeting shRNA (Sigma SHC216) was used as a control.

### Lonafarnib and Methylstat Treatment

HUVECs were treated with either 1.0 μM Methylstat (Sigma-Aldrich, SML0343), a histone demethyltransferase inhibitor or 0.5μM Lonafarnib (Tocris, 6265), a farnesyltransferase inhibitor. HUVECs were treated with a daily dose of 1.0 μM Methylstat (Sigma-Aldrich, SML0343) for 48 hours. HUVECs were treated with a daily dose of 0.5μM Lonafarnib (Tocris, 6265) for 72 hours.

### Fluid Shear Stress

HUVECs were seeded onto ibidi chamber slides (ibidi-treated μ-slides I^0.6^, cat #80186 or ibidi-treated μ-slides VI^0.4^, cat#80606, Germany), coated with 60ng/mL fibronectin (Sigma-Aldrich, F1141). At 80% confluency, HUVECs were exposed to laminar or oscillatory shear stress using the ibidi pump system (ibidi, cat #10902, Germany), at 12 dynes/cm^2^ for 1, 3, or 6 days with or without modifications perfused in the media.

### Cell Fixation and Labelling

After fluid shear stress experiments were finished, HUVECs were washed two times with PBS and fixed for 10 min at room temperature with 4% paraformaldehyde in PBS. After three washes with PBS, the cells were permeabilized for 10 min at room temperature with 0.2% Triton X-100 in PBS and blocked with 5% BSA for 1 hour at room temperature. Cells were then incubated overnight at 4 degrees Celsius room temperature with the primary Ab diluted in blocking solution, using either anti-lamin A antibody (cat # sc-7292, Santa Cruz Biotechnology) for control cells, or anti-HA antibody (cat # 901501, Biolegend) for progerin-expressing cells and anti-lamin B1 antibody (cat # ab16048, Abcam). Three more washes with PBS were then followed by incubation with the secondary Ab (Alexa Fluor 647-conjugated donkey anti-mouse IgG; Thermo Fisher) and stained with rhodamine phalloidin (cat # PHDR1, Cytoskeleton) for 45 min followed by three additional PBS washes. Samples were stained with Hoechst 33342 (ThermoFisher) and mounted with ibidi mounting medium (ibidi, cat #50001, Germany). For cell death assays, an apoptosis/necrosis Detection kit (abcam, ab176749) was used following the manufacturer’s instructions.

### Quantification and Imaging Analysis

Samples were imaged on Zeiss LSM 710 confocal microscope at 20x and 63x. Image analysis was completed using Fuji Image J. We analyzed the progerin-expressing endothelial cell response to the force of shear stress by quantifying abnormally structured nuclear lamina (through lamin A, HA-progerin, and lamin B staining), nuclear lamina outward blebs (through lamin A and HA-progerin staining), and the presence or absence of micronuclei (through Hoechst staining). Cell loss was quantified using a cell counting macros on Image J.

### Statistical Analysis

Statistical significance was measured using an unpaired, two-tailed student t-test for data containing two groups. For data involving more than two groups, the Analysis of Variance (ANOVA) test was performed in order to obtain the statistical analysis for the data sets concerned. A further comparison of the groups was performed using the Tukey (HSD) test to determine significant differences between groups. All statistical tests were conducted at a 5% significance level (p<0.05). Prism GraphPad was used for statistical analyses.

## Results

### Endothelial cells expressing progerin or overexpressing wild-type lamin A are unable to adapt to shear stress

Previous work has shown that HGPS cells exhibit both a combination of a stiffened nucleoskeleton and a softened nuclear interior,(14) which in turn can cause mechanical irregularities and impaired mechanoadapatation (14, 34). We therefore sought to understand how progerin expression would affect endothelial cell responses to fluid shear stress, the frictional drag force created by blood flow. We expressed progerin in HUVECs using an adenovirus, and as a control we also examined the effects of overexpression (OE) of wild-type lamin A. HUVECs expressing either progerin or overexpressing wild-type lamin A experienced dramatic cell loss after 72 hours of arterial levels of shear stress (12 dynes/cm^2^) [**Figure 1 (A & B)**]. Interestingly there was minimal cell loss for both groups at 24 hours of shear stress, indicating that cell loss is not a rapid event. Non-transuded control cells had no cell loss and were able to characteristically align in the direction of shear stress. Additionally, no cell loss was observed for progerin-expressing or lamin A OE HUVECs when grown in static culture (0-hour images), indicating that fluid shear stress is necessary for cell loss. Additionally, an apoptosis assay showed progerin-expressing HUVECs under 24 hours of laminar shear stress have increased surface expression of the apoptotic marker phosphatidyslerine **[Supplemental Figure 1]**.

**Figure 1:**
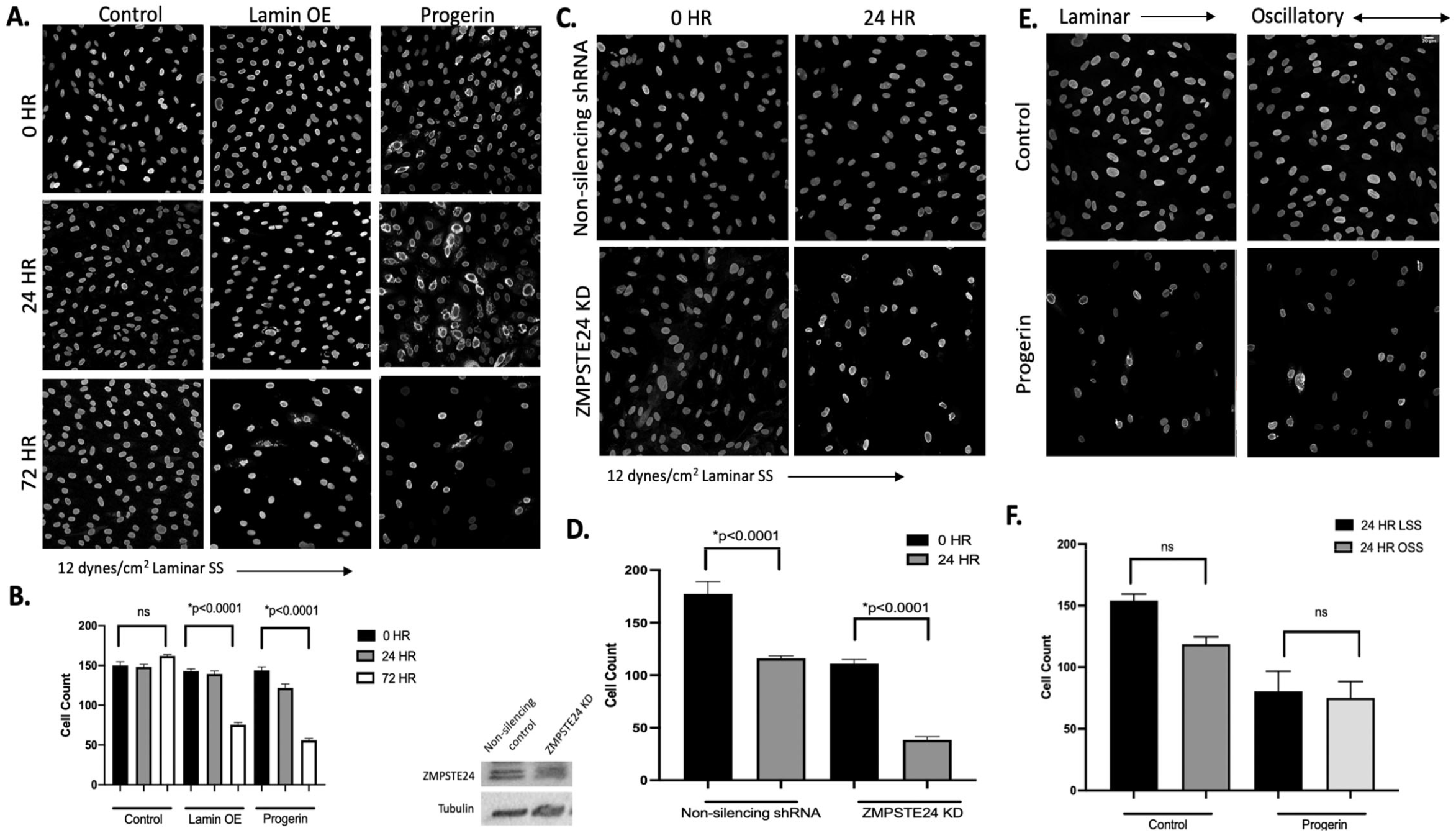
Progerin expression causes cell loss in HUVECs under laminar shear stress. Confocal images taken at 20x. HUVECs progerin (HA stained). Lamin OE, Control, ZMPSTE24 KD, and non-silencing shRNA (lamin A/C stained). Ordinary One-way ANOVA. N=3. (**A**) Cell loss in progerin-expressing HUVECs under laminar shear stress for 0, 24 and 72 hours. (**B**) Cell count for HUVECs under laminar shear stress for 0, 24, and 72 hours. (**C**) Knockdown of the ZMPSTE24 gene with RNAi leads to a farnesylated form of lamin A. Cell loss in ZMPSTE24 under laminar shear stress for 0 and 24 hours was observed. (**D**) Cell count for ZMPSTE24 KD under laminar shear stress for 0 and 24 hours. (**E, F**) No difference was observed between 24 hours of oscillatory and laminar shear forces in progerin-expressing HUVECs.

As an alternate approach to model HGPS in endothelial cells we developed HUVECs in which ZMPSTE24 was knocked down by shRNA. ZMPSTE24 is a protease involved in post-translational cleavage of farnesylated prelamin A. Loss of ZMPSTE24 leads to the accumulation of farnesylated lamin A and phenocopies features of HGPS (11, 12). HUVECs with knockdown of ZMPSTE24 also experienced similar cell loss when exposed to shear stress, [**Figure 1 (C & D)**]. To see if oscillatory flow would cause more cell death events in the progerin-expressing cells we exposed the HUVECs to 12 dynes/cm^2^ of laminar and oscillatory shear stress. No significant difference in cell loss was observed between the different flow types [**Figure 1 (E & F)**]. Taken together, these results show that changes in nuclear lamins can affect the ability of endothelial cells to adapt to the forces of fluid flow.

### Progerin-expressing endothelial cells have increased nuclear abnormalities in both static and shear conditions

Progerin-expressing HUVECs under both flow **[Figure 1(A)]** and static conditions displayed exaggerated nuclear morphologies. Previous work as shown that the lamina in HGPS cells has a significantly reduced ability to rearrange under mechanical stress (13). We quantified nuclear abnormalities (including dysmorphic nuclear lamina, outward blebs, and micronuclei) in progerin-expressing HUVECs under shear stress and static conditions, and compared results to control and the lamin OE groups. Consistent with prior observations in non-endothelial cell types, nuclear abnormalities were present in progerin-expressing endothelial cells. **Figure 2(A)** shows the examples of dysmorphic nuclear lamina in both progerin-expressing and lamin OE cells. In the progerin-expressing cells we also observed disruptions in lamin B structure. We also examined the incidence of outward blebs **[Figure 2 (B)]** and micronuclei **[Figure 2 (C)]**. Quantification of dysmorphic nuclear lamina, outward blebs, and micronuclei showed similar levels of occurrence both under shear stress **[Figure 2 (D)]** and static culture conditions **[Figure 2 (E)]** for progerin expressing cells.

**Figure 2:**
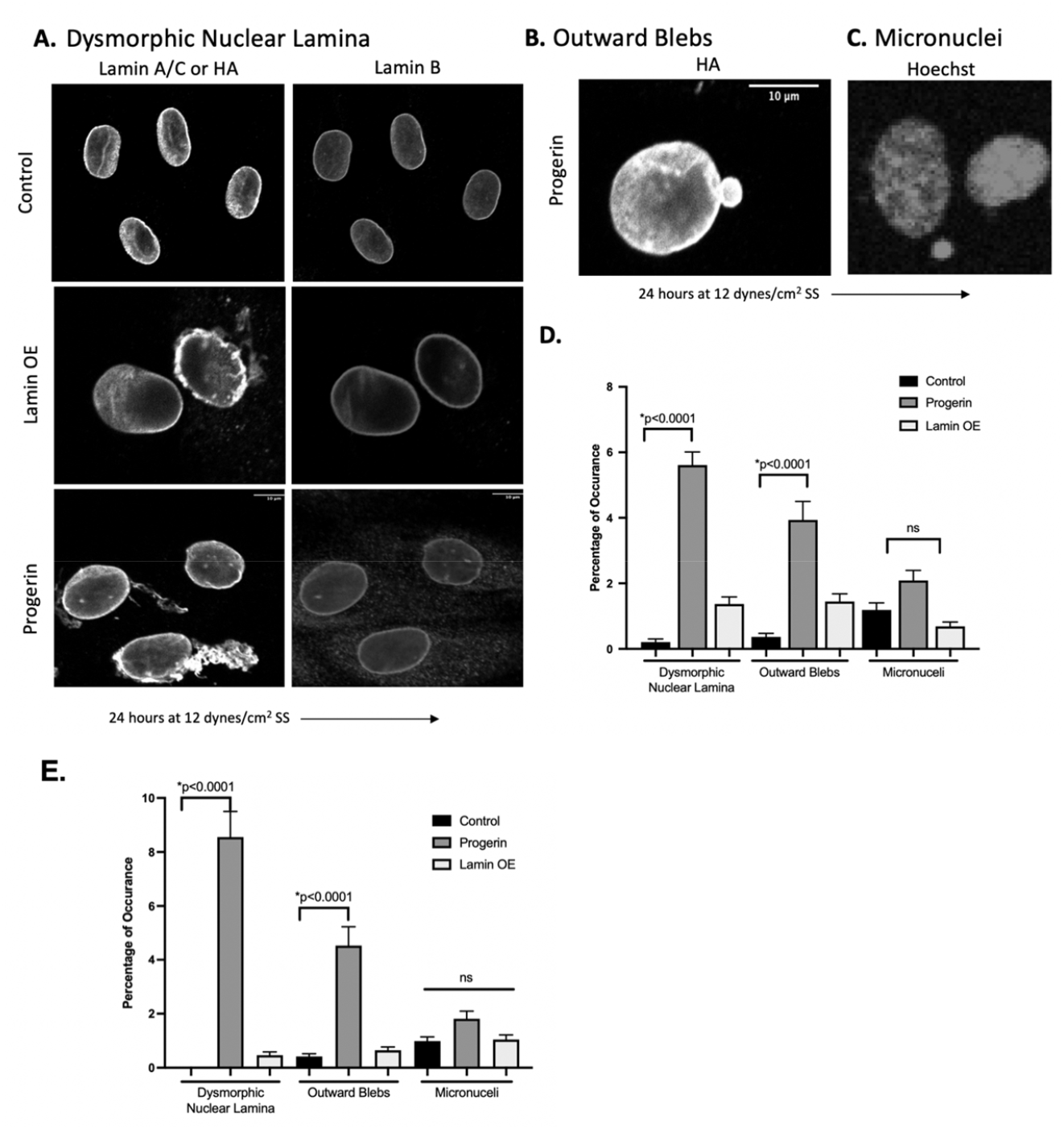
Laminar shear stress causes nuclear envelope disruption in Progerin-expressing HUVECs. Confocal images taken at 20x, 40x, and 63x. HUVECs HA-progerin (HA and lamin B stained), and Control (lamin A/C and lamin B stained). Ordinary one-way ANOVA. N=3.The rigidity of the progerin nuclei in HUVECs have shown an inability to properly regulate physiological force. **(A)** Dysmorphic nuclear lamina characterized by abnormal structure HA-progerin and lamin B staining in progerin-expressing HUVECs under 24 hours of laminar shear stress. **(B & C)** lamin A/C blebs and micronuclei was observed in progerin-expressing cells after 24 hours of shear stress. **(D)** Quantification of nuclear abnormalities in shear stress condition. **(E)** Quantification of nuclear abnormalities in static condition.

### Lonafarnib rescues cell loss and nuclear envelope dysmorphia in progerin-expressing HUVECs under shear stress

In progerin-expressing cells, the attachment of farnesyl groups causes the nuclear envelope to have lobes instead of a round shape. This lobulation of the nuclear envelope is due to accumulation of progerin within the nucleus and dramatically changes the nuclear architecture as well as its stability (21). Previously it has been shown that the farnesyltransferase inhibitor (FTI) Lonafarnib can be used to prevent progerin accumulation and improve nuclear shape (35), and is currently used in clinical trials as a treatment for HGPS. To inhibit farnesylation of lamin A, ZMPSTE24 KD and progerin-expressing HUVECs were treated with Lonafarnib for 72 hours. These treated cells were then subjected to 24 hours of laminar shear stress. We hypothesized that inhibiting progerin farnesylation would improve the ability of these cells to adapt to shear stress. Our results showed that Lonafarnib significantly prevented cell loss the ZMPSTE24 KD cells [**Figure 3 (A & B)**] but did not lead to a significant improvement in the progerin-expressing cells. Furthermore, the results showed that Lonafarnib treatment prevented nuclear envelope disruptions in the HA-progerin expressing cells **[Figure 3 (C & D)]**. Taken together, these improvements directly demonstrate the ability of Lonafarnib to improve endothelial nuclear morphology in progerin-expressing cells, as well as enhancing the ability of these cells to adapt to changes in mechanical forces.

**Figure 3:**
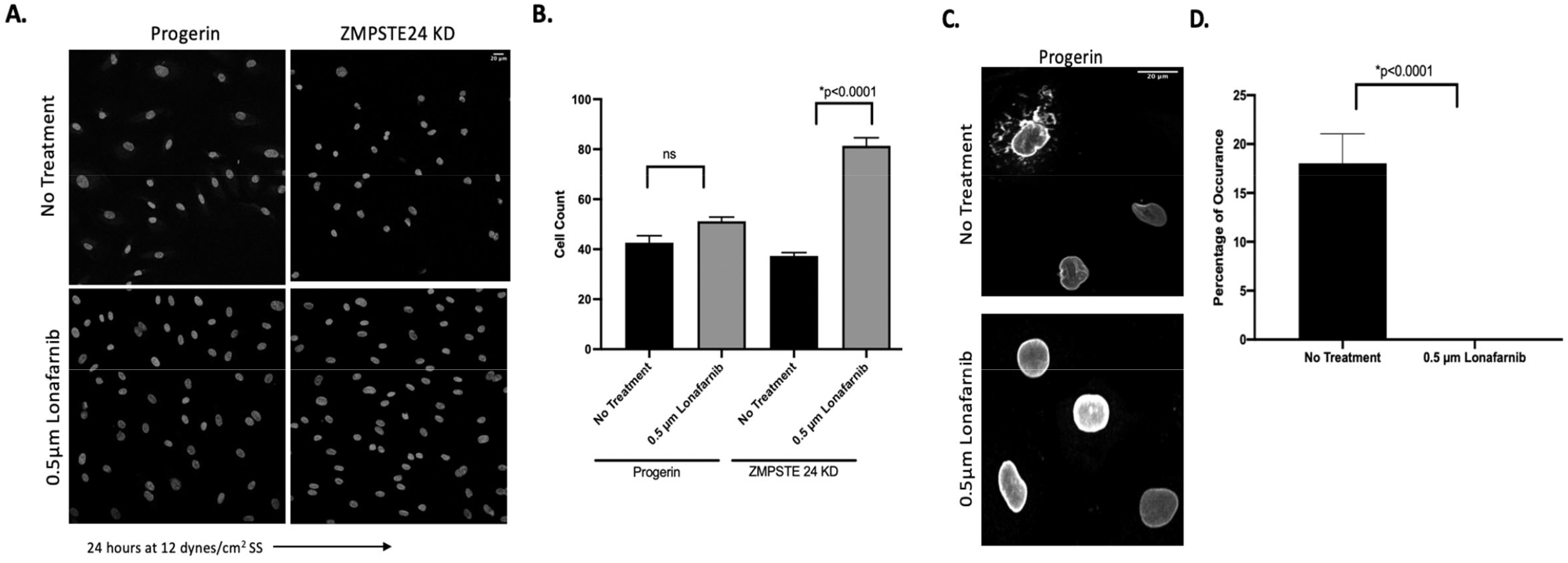
Lonafarnib rescues cell loss and nuclear envelope disruptions in Progerin-expressing HUVECs under shear stress. HUVECs were treated with a daily dose of 0.5μM Lonafarnib (Tocris, 6265) for 72 hours and then exposed to fluid flow. **(A & B)** Lonafarnib rescues cell loss in ZMPSTE24 KD cells under 24 hours of shear stress. Confocal images taken at 20x. HUVECs stained with Hoechst. Ordinary One-way ANOVA. N=3. **(C & D)** Lonafarnib rescues nuclear envelope disruptions in progerin-expressing HUVECs under shear stress. Confocal images taken at 40x. HUVECs stained with HA-progerin. Unpaired T-test. N=3.

### Methylstat recuses cell loss in progerin-expressing HUVECs under shear stress

It has been previously shown that progerin-expressing cells have alternations in histone modifications, including: a loss of peripheral heterochromatin, reduced levels of H3k9me3 and increased levels of trimethylation of H4K20, an epigenetic mark for constitutive heterochromatin on H4 (17). Notably, pharmacological-induced increases in heterochromatin have been shown to rescue nuclear morphology in a HGPS patient cells (36). To examine if increases in heterochromatin would improve the ability of progerin-expressing endothelial cells to adapt to shear stress, we used the drug methylstat, an inhibitor of histone trimethyl demethylases. ZMPSTE24 KD and progerin-expressing HUVECs were treated with Methylstat for 72 hours and exposed to 24 hours of laminar shear stress. Cell loss **[Figure 4 (A & B)]** and nuclear envelope disruptions were prevented [**Figure 4 (C & D)**] in cells treated with Methylstat. Taken together, these results show that increases in DNA methylation in progerin-expressing endothelial cells rescue nuclear morphology and ability to adapt to shear stress.

**Figure 4:**
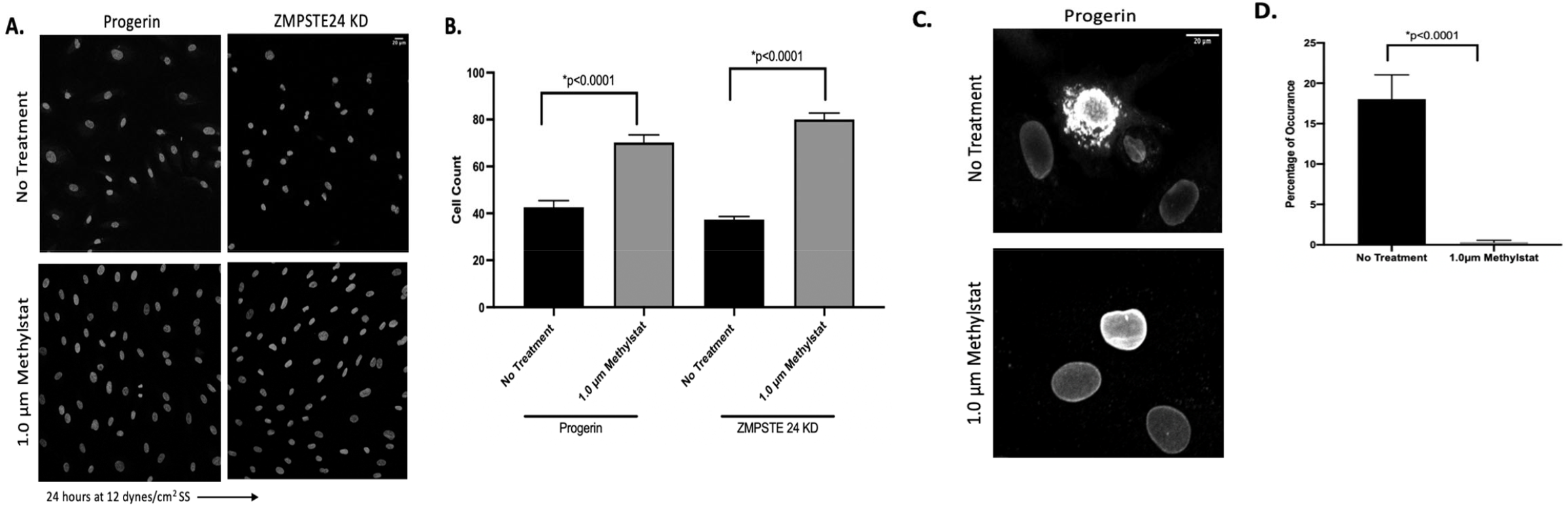
Methylstat recuses cell loss in Progerin-expressing HUVECs under shear stress. HUVECs were treated with a daily dose of 1.0μM Methylstat (Sigma-Aldrich, SML0343) for 48 hours and then exposed to fluid flow. **(A & B)** Methylstat rescues cell loss in progerin-expressing HUVECs under shear stress. Confocal images taken at 20x. HUVECs stained with Hoechst. Ordinary One-way ANOVA. N=3. **(C & D)** Methylstat rescues nuclear envelope disruptions in progerin-expressing HUVECs under shear stress. Confocal images taken at 40x. HUVECs stained with HA-progerin. Unpaired T-test. N=3.

### Pre-alignment of progerin-Expressing HUVECs prevents cell loss under shear stress

We hypothesized that progerin-expressing endothelial cells which did not have to undergo cellular and nuclear shape changes would be less affected by exposure to shear stress. To investigate how progerin expression would affect aligned endothelial cells, we first exposed nontransduced HUVECs to shear stress for 72 hours to induce alignment. Afterwards, these cells were then transduced with progerin or a control adenovirus (GFP), and exposed to an additional 72 hours of shear stress. Our results showed that progerin expression in pre-aligned cells [**Figure 5 (A)**] resulted in reduced cell loss **[Figure 5 (C)]** and nuclear envelope disruptions **[Figure 5 (D)]** when compared to progerin-expressing cells not adapted to shear stress [**Figure 5 (B)**].

**Figure 5:**
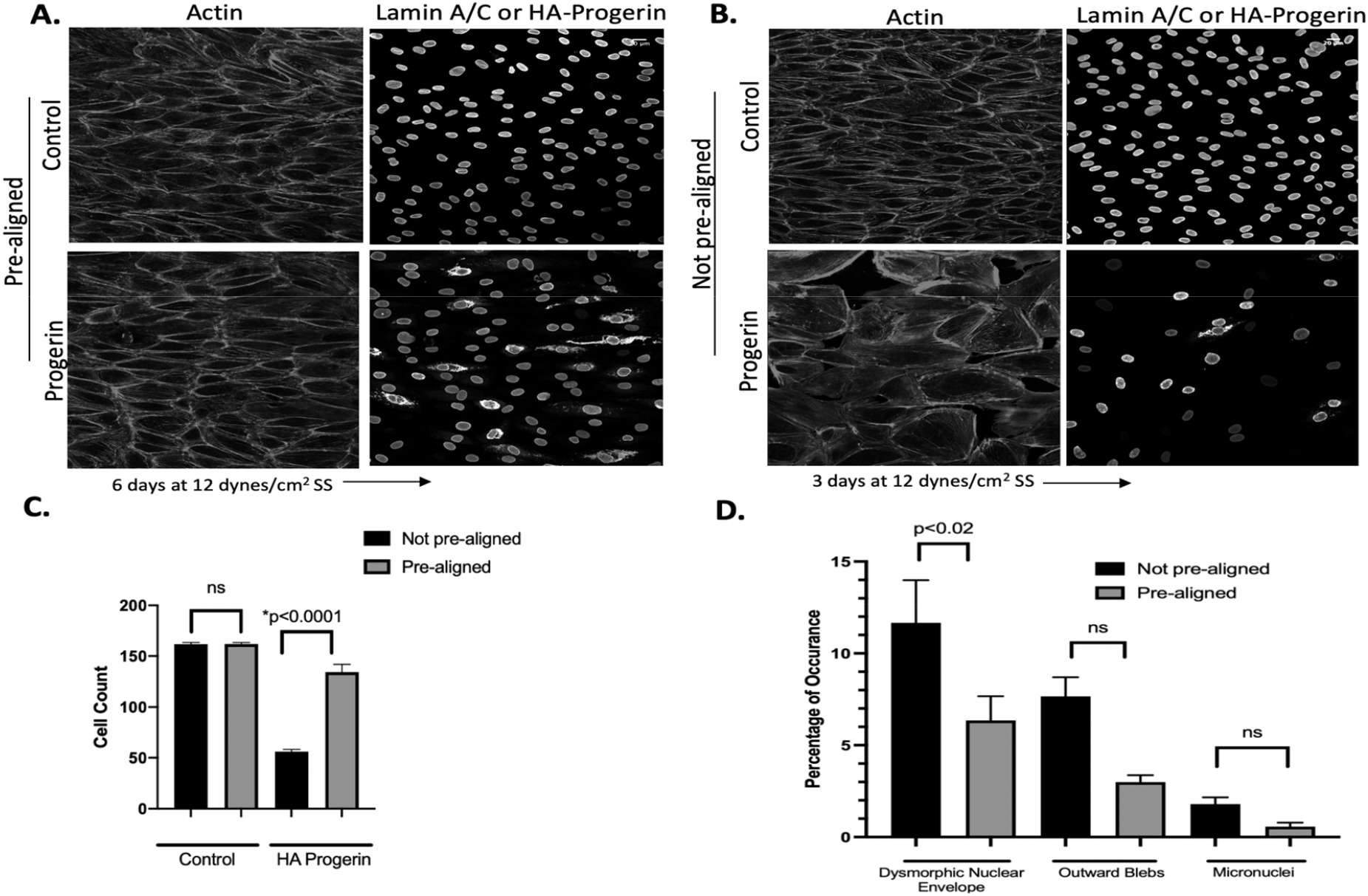
Pre-alignment of Progerin-Expressing HUVECS prevents cell loss under shear stress. Confocal images taken at 20x. HUVECs stained with HA-progerin and Control stained with lamin A/C. Both groups stained with Phalloidin. One-way ANOVA. N=3. Progerin was expressed in both pre-aligned (A) and not pre-aligned cells (B) and exposed to 72 hours of shear stress at 12 dynes/cm2. Pre-aligned cells had reduced cell loss (C) and nuclear envelope disruptions (D) when progerin was expressed after actin alignment.

## Discussion

In this work we developed an *in vitro* model of HGPS endothelium by 1) expressing progerin and 2) knockdown of ZMPSTE24 in HUVEC. In both instances these cells exhibited characteristic progerin-induced changes in nuclear morphology, indicating that the endothelial cell nucleus is sensitive to progerin expression. Strikingly, application of fluid shear stress resulted in dramatic cell loss, occurring between 1-3 days of fluid shear stress **[Figure 1]**. Interestingly, pre-alignment of endothelial cells to fluid shear stress prior to expression of progerin **[Figure 5]** prevented the cell loss, suggesting that progerin-expression prevents the ability of endothelial cells to adapt to changes in mechanical forces, but has less of an effect in cells already adapted and aligned to shear stress.

Although a number of prior studies have focused on the effects of progerin in fibroblasts and vascular smooth muscle cells(20, 24, 29, 30, 37–39), our results add to growing evidence that the endothelium is also sensitive to progerin expression (31, 32, 40). Although two prior mouse models expressing progerin in the endothelium did not report dramatic loss of endothelial cells (31, 32), our finding that pre-aligned cells are less sensitive to progerin expression suggests that progerin expression may be most significant when endothelial cells are required to adapt to changes in mechanical environments. Supporting this hypothesis *Sun et al* showed that there was defective neovascularization of progerin-expressing endothelial cells in response to ischemia(31). Additionally, *Osmanagic-Myers et al* showed impaired endothelial alignment in response to short-term (3 hours) fluid shear stress (32). Loss of endothelial cells is especially significant in the context of HGPS early onset atherosclerosis, as endothelial dysfunction and damage is considered an initial step in the onset of atherosclerosis(41).

We showed that two pharmacological treatments previously shown to improve HGPS nuclear morphology, lonafarnib (farnesyltransferase inhibitor) (35) and methylstat (inhibitor of histone trimethyl demethylases) (36) also restored normal nuclear morphology to progerin-expressing endothelial cells **[Figures 3 and 4]**. Interestingly increasing chromatin methylation with methylstat significantly improved the ability of progerin-expressing endothelial cells to adapt to fluid shear stress **[Figure 4]**. However, lonafarnib did not rescue cell loss for progerin-expressing cells (but did rescue ZMPSTE24 knockdown cells). Thus, improvements in nuclear morphology did not uniformly result in improved mechanoadaptation to shear stress.

An unanswered question in our work is the specific biochemical or physical mechanisms that account for the loss of progerin-expressing cells under fluid shear stress. Our group recently published work showing that the LINC complex is a necessary structure in endothelial cell mechanoadapation, in which we observed that disruption of the LINC complex resulted in a much more rapid loss of cells under fluid shear stress, due in part to impaired cell-substrate attachment (42). We do not believe the progerin-expressing cells have weaker or impaired attachment to the substrate, as we observed no cell loss or detachment in static culture, as well as minimal cell loss after one day of shear stress. In progerin-expressing endothelial cells we did observe an increase in expression of surface phosphatidylserine **[Supplemental Figure 1]**, suggesting that altered mechanical or biochemical signaling in progerin-expressing leads to cell loss through apoptosis. Related, prior work by *Bidault et al* showed progerin-expressing in endothelial cells have increased markers of DNA damage as well as upregulation of p53 and p21 which induce cellular senescence(40).

Interestingly, we observed that similar levels of cells were lost under shear stress when wildtype lamin A was overexpressed as compared to progerin expression **[Figure 1A]**. However, lamin A overexpressing cells had significantly less dysmorphic nuclear lamina and blebbing when compared to progerin expressing cells **[Figure 2D]**. Thus, changes in nuclear morphology do not completely account for the cell loss or an inability to adapt to shear stress. Overexpression of lamin A has been shown to increase nuclear stiffness(43), similar to progerin expression(14). It is therefore tempting to speculate that stiffer nuclei are less able to adapt to changing mechanical forces. Prior work has shown that there are substantial changes in nuclear shape and stiffness when endothelial cells adapt to shear stress (44), indicating that the nucleus undergoes significant remodeling to adapt to shear stress.

This work highlights the nuclear lamina as a critical feature for endothelial adaptability to fluid shear stress. We hypothesize that other factors that control nuclear stiffness beyond nuclear lamins, such as chromatin stiffness, may also impact how readily endothelial cells adapt to shear stress. We also note that aging can induce similar nuclear phenotypes to HGPS, including expression of progerin, increased nuclear stiffness, and altered nuclear morphology(45). An important question will be to determine if aging-associated changes in nuclear stiffness similarly impair endothelial cell adaptation to mechanical forces.

## Acknowledgements

We acknowledge Andrew Stephens for helpful discussions. This work was supported by a NSF Graduate Research Fellowship (to BD), NIH grant R35 GM119617 (to DC), and NSF CAREER award CMMI 1653299 (to DC). Services in support of the research project were generated by the VCU Microscopy Shared Resource, in part, with funding from NIH-NCI Cancer Center support grant P30 CA016059.

## Author Contributions

B.D., K.D., and D.C. designed the study. N.N, L.S., K.B., H.P. and B.D. performed experiments. H.P. and B.D. analyzed collected data. B.D. and D.C. wrote the manuscript.

---

**Supplemental Figure 1: Progerin-expressing cells undergo apoptosis under shear stress.** Images taken with confocal microscope at 20x. A cell death assay showed progerin-expressing HUVECs under 24 hours of laminar shear stress experience apoptosis, indicated by the presence of the Phosphatidyslerine, an apoptotic marker.

