## Supplementary figures and images for "Progerin-Expressing Endothelial Cells are Unable to Adapt to Shear Stress"

### Supplemental Figure 1

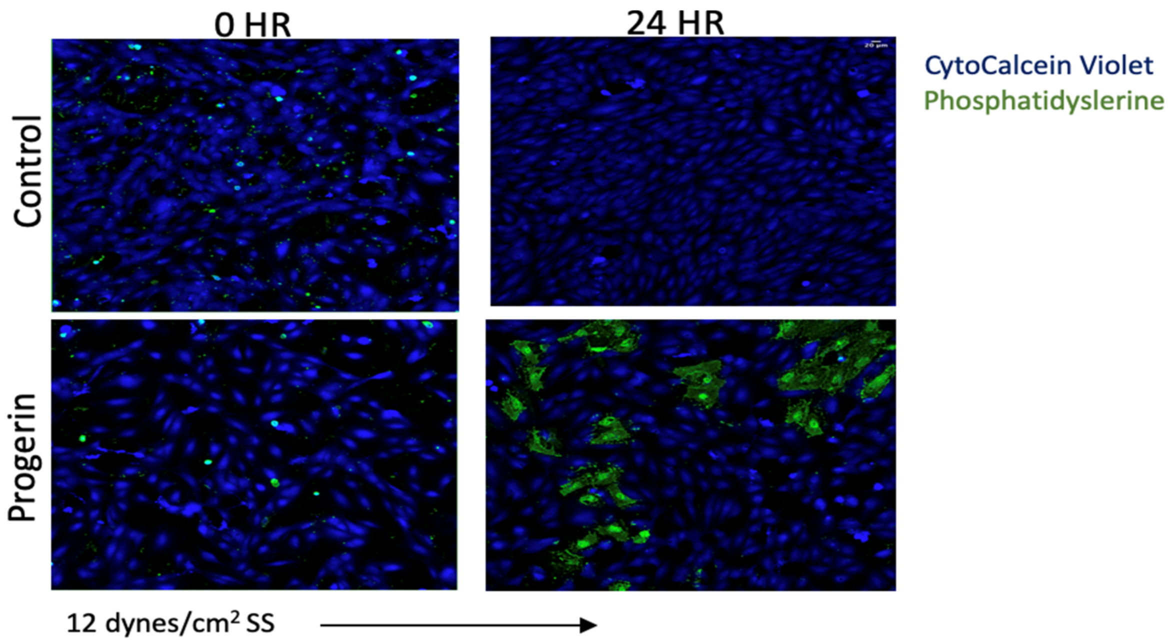
